# Visualizing the itch-sensing skin arbors

**DOI:** 10.1101/2020.05.18.098871

**Authors:** Yanyan Xing, Haley R. Steel, Henry B. Hilley, Katy Lawson, Taylor Niehoff, Liang Han

**Author notes:** These authors contributed equally to the manuscript (Yanyan Xing, Haley R. Steel). Corresponding author: Liang Han, PhD, School of Biological Sciences, Georgia Institute of Technology, Atlanta, GA 30332, USA.

## Abstract

Diverse sensory neurons exhibit distinct neuronal morphologies with a variety of axon terminal arborizations used to subserve their functions. Due to its clinical significance, the molecular and cellular mechanisms of itch are being intensely studied. However, a complete analysis of itch-sensing terminal arborization morphology is missing. Using a novel *MrgprC11^CreERT2^* transgenic mouse line, we labeled a small subset of itch-sensing neurons that express multiple itch-related molecules including MrgprA3, MrgprC11, histamine receptor H1, IL-31 receptor, 5-HT receptor 1F, natriuretic precursor peptide B, and neuromedin B. By combining sparse genetic labeling and whole-mount PLAP histochemistry, we found that itch-sensing skin arbors exhibit free endings with extensive axonal branching in the superficial epidermis and large receptive fields. These results revealed the unique morphological characteristics of itch-sensing neurons and provide novel insights into the basic mechanisms of itch transmission.

## INTRODUCTION

Primary somatosensory neurons detect nociception, mechanical stimulation, and body location via peripheral axons, located in both the skin and the body interior, and transmit signals to the spinal cord via central axons. Diverse sensory neurons are classified by molecular composition, neurophysiological properties (axon diameter, myelination, and conduction velocity), and defined functions. This diversity is also evidenced by distinct innervation patterns involving the morphology of peripheral and central axon terminal arborizations, specialized peripheral anatomical structures associated, termination zones within particular lamina of the spinal cord dorsal horn, and somatotopic organization (Abraira and Ginty, 2013; Basbaum et al., 2009; Crawford and Caterina, 2020; Ikoma et al., 2006). For example, most cutaneous mechanosensors innervate hair follicles and skin touch organs, such as Merkel cells and Meissner corpuscle, and project central axons to the laminar III-V of the spinal cord dorsal horn (Abraira and Ginty, 2013). In contrast, nociceptors mediating pain and itch innervate the epidermis as free nerve endings and their central axons terminate within the outermost lamina of the dorsal horn (lamina I and II) (Han et al., 2013; Zylka et al., 2005). Thus, morphological and anatomical analysis of sensory innervation patterns is fundamental to understand somatosensory processing. Indeed, deep functional insights on touch sensation and sensory acuity were revealed from the integration of morphological and physiological characterizations of several subtypes of sensory neurons including the low-threshold mechanoreceptors (LTMRs) and non-peptidergic nociceptors (Bai et al., 2015; Ghitani et al., 2017; Kuehn et al., 2019; Li and Ginty, 2014; Li et al., 2011; Olson et al., 2017; Rutlin et al., 2014).

Itch, a somatosensory modality mainly arising from the skin, serves as an alert to potential threats. However, chronic itch associated with dermatological and systemic disorders is a debilitating symptom that is often refractory to available treatments (Ikoma et al., 2006). Three distinct itch-sensing neuronal populations were previously identified by their unique combination of itch receptors: MrgprA3^+^ neurons are enriched with MrgprC11 and histamine receptors (Han et al., 2013; Liu et al., 2009; Xing et al., 2020); Nppb^+^ neurons express receptors for IL31 (Il31ra), cysteinyl leukotriene (Cysltr2), serotonin (Htr1f), and sphingosine-1-phosphate (S1pr1) (Solinski et al., 2019); and MrgprD^+^ neurons express lysophosphatidic acid receptors (Lpar3 and Lpar5) (Sharma et al., 2020; Usoskin et al., 2015). While recent investigations have given us a glimpse of the molecular, cellular, and circuitry basis of itch processing, analysis of the innervation pattern of itch-sensing neurons is missing.

Detailed morphological and anatomical analysis of sensory neurons requires visualization of single sensory arbor, which is technically challenging because sensory arbors always overlap and intermingle. In the present study, by combining spare genetic labeling and whole-mount PLAP histochemistry staining we successfully trace the morphology of a subset of itch-sensing neurons. These neurons are mainly peptidergic and express multiple itch-related molecules including MrgprA3, MrgprC11, IL31ra, Hrh1, Htr1f, Nppb, and Nmb. Peripheral itch-sensing arbors in the skin are characterized by free endings with extensive axonal branching in the superficial epidermis. Their arbor areas are much larger than the non-itch-sensing MrgprB4^+^ and MrgprD^+^ arbors, suggesting that itch-sensing neurons have large receptive fields. These results reveal the unique morphology of itch-sensing arbors and help us to understand how itch is initiated in the skin.

## RESULTS

### MrgprC11^CreERT2^ leaky neurons label a small subset of peptidergic nociceptors

MrgprC11 is a recently identified itch receptor in the Mas-related G protein-couple receptor (Mrgpr) family. It is specifically expressed in small-diameter DRG sensory neurons with several identified pruritogen agonists including Bam8-22, SLIGRL, and Cathepsin S (Liu et al., 2009; Liu et al., 2011; Reddy et al., 2015). To investigate the morphological characteristics of itch-sensing neurons, we generated a BAC transgenic *MrgprC11^CreERT2^* mouse line and utilized the tamoxifen-inducible CreERT2/loxP system to perform sparse genetic labeling and neuronal morphology tracing. Crossing *MrgprC11^CreERT2^* mice with the Cre-dependent *Rosa26^tdtomato^* line yielded sparse recombination in 0.94% of DRG sensory neurons (45 out of 4772 cells, referred to as MrgprC11^CreERT2^ leaky cells) in the absence of tamoxifen treatment (Figure 1). These sparsely labeled DRG neurons are small diameter (260.11 ± 14.82 μm^2^) neurons positive for MrgprC11 (95%, Figure 1A), the nociceptive marker TRPV1 (94%, Figure 1B), and the peptidergic marker substance P (93%, Figure 1C), and negative for myelinated neuron marker neurofilament 200kD (NF200, Figure 1G). A subset were positive for peptidergic marker calcitonin gene-related peptide (CGRP, 54%, Figure 1D), the nonpeptidergic nociceptive marker IB4 (17%, Figure 1E), and the purinergic receptor X3 (P2X3, 31%, Figure 1F). Moreover, these neurons are distinct from the MrgprB4^+^ neurons (Figure 1H) whose arbor morphology were previously revealed (Liu et al., 2007). Therefore, MrgprC11^CreERT2^ leaky neurons are a small subset of peptidergic nociceptors.

**Figure 1.**
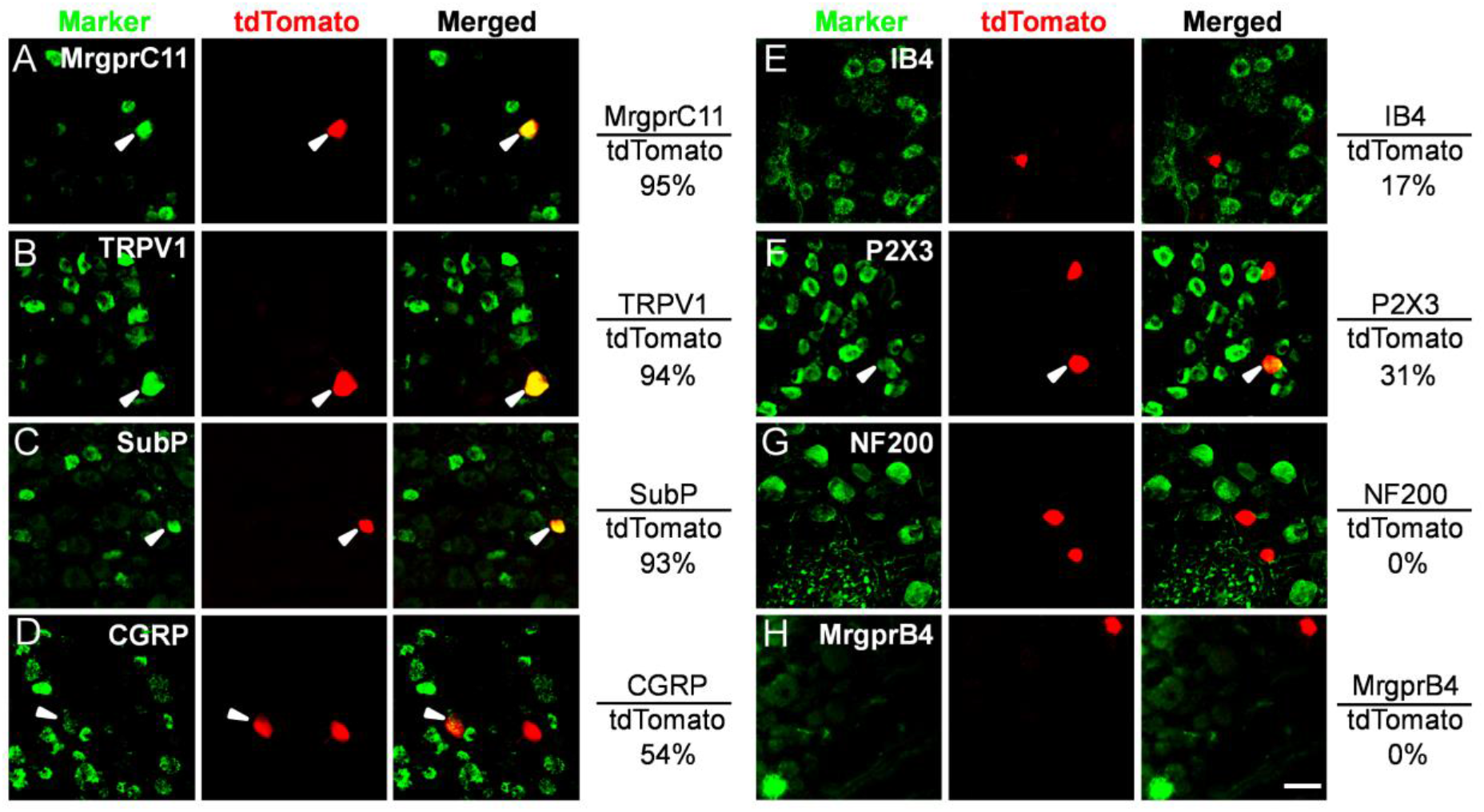
MrgprC11^CreERT2^ leaky neurons label a small subset of peptidergic nociceptors. (A-H) Double immunostaining of lumbar level 3-5 (L3-5) DRG sections from *MrgprC11^tdTomato^* mice to detect indicated sensory neuron markers in *MrgprC11^CreERT2^* leaky neurons. Overlap quantifications were shown on the right of each panel (% of tdTomato^+^ cells that express markers,). Arrowheads point to overlapping cells. Scale bars = 50 μm. N = 3 mice for each marker.

### MrgprC11^CreERT2^ leaky neurons represent itch-sensing neurons expressing multiple itch receptors

Surprisingly, MrgprC11^CreERT2^ leaky cells expressed multiple itch-related molecules including MrgprA3 (60%, Figure 2A) and Nppb (70%, Figure 2C), markers for two distinct itch-sensing subtypes (Han et al., 2013; Sharma et al., 2020; Solinski et al., 2019; Usoskin et al., 2015). The majority of neurons expressed histamine receptor (Hrh1, 73.3%, Figure 2B), which are enriched in MrgprA3^+^ neurons. They were also positive for Il31ra (76.9%, Figure 2D) and Htrf1 (50%, Figure 2E), which are enriched in Nppb^+^ neurons. Moreover, 62.1% of them expressed neuromedin B (Nmb, Figure 2F), a neuropeptide required for histaminergic itch (Wan et al., 2017). Very few of them expressed MrgprD or Lpar3 (3.7% and 4.2% respectively, Figure 2G-H), markers for MrgprD^+^ itch-sensing neurons. The expression of multiple itch-related molecules in MrgprC11^CreERT2^ leaky cells indicates that they are itch-sensing neurons.

**Figure 2.**
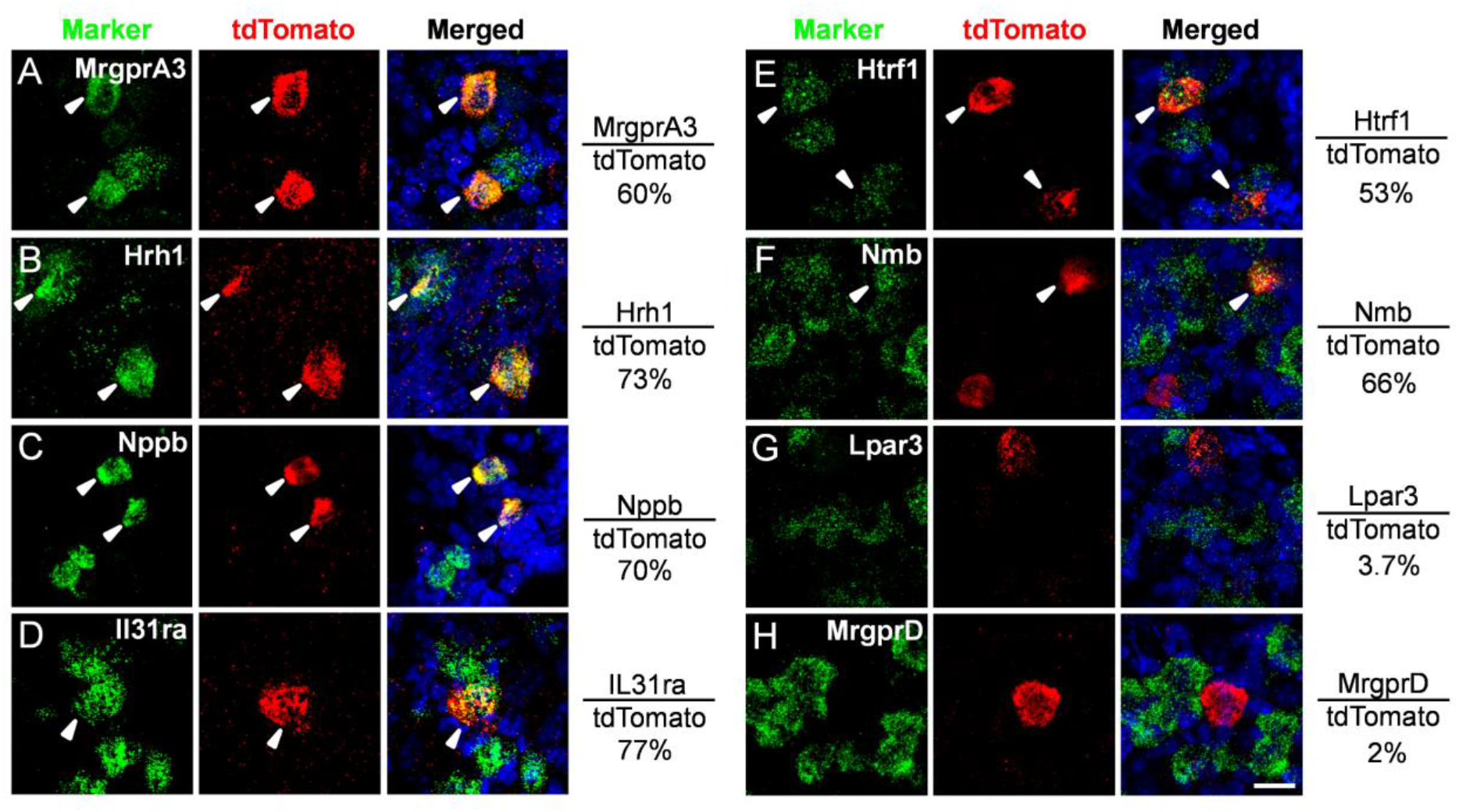
MrgprC11^CreERT2^ leaky neurons express multiple itch receptors. (A-H) RNAscope fluorescent *in situ* hybridization detecting itch-related molecules using lumbar DRG sections from *MrgprC11^tdTomato^* mice. Overlap quantifications were shown on the right of each panel (% of tdTomato^+^ cells that express markers). Arrowheads point to overlapping cells. Scale bars = 25 μm. N = 3 mice for each marker.

We next performed calcium imaging with cultured dissociated neurons to examine if they exhibit neuronal responses to pruritogens. We generated *MrgprC11^tdTomato^*; *Pirt^GCaMP3^* mice that express the calcium indicator GCaMP3 in all sensory neurons (Kim et al., 2014). MrgprC11^CreERT2^ leaky cells were labeled by tdTomato fluorescence. Consistent with the expression of itch receptors, most MrgprC11^CreERT2^ leaky neurons responded to MrgprA3 agonist chloroquine (90%), MrgprC11 agonist Bam8-22 (91.7%), histamine (88.3%), IL-31 (84.9%), and 5-HT1F receptor agonist LY344864 (65.3%). Almost all cells also responded to the TRPV1 agonist capsaicin (97.9%) (Figure 3A-B and 3D). None of the tested cells responded to the application of MrgprD agonist β-alanine or LPA (Figure 3C-D).

**Figure 3.**
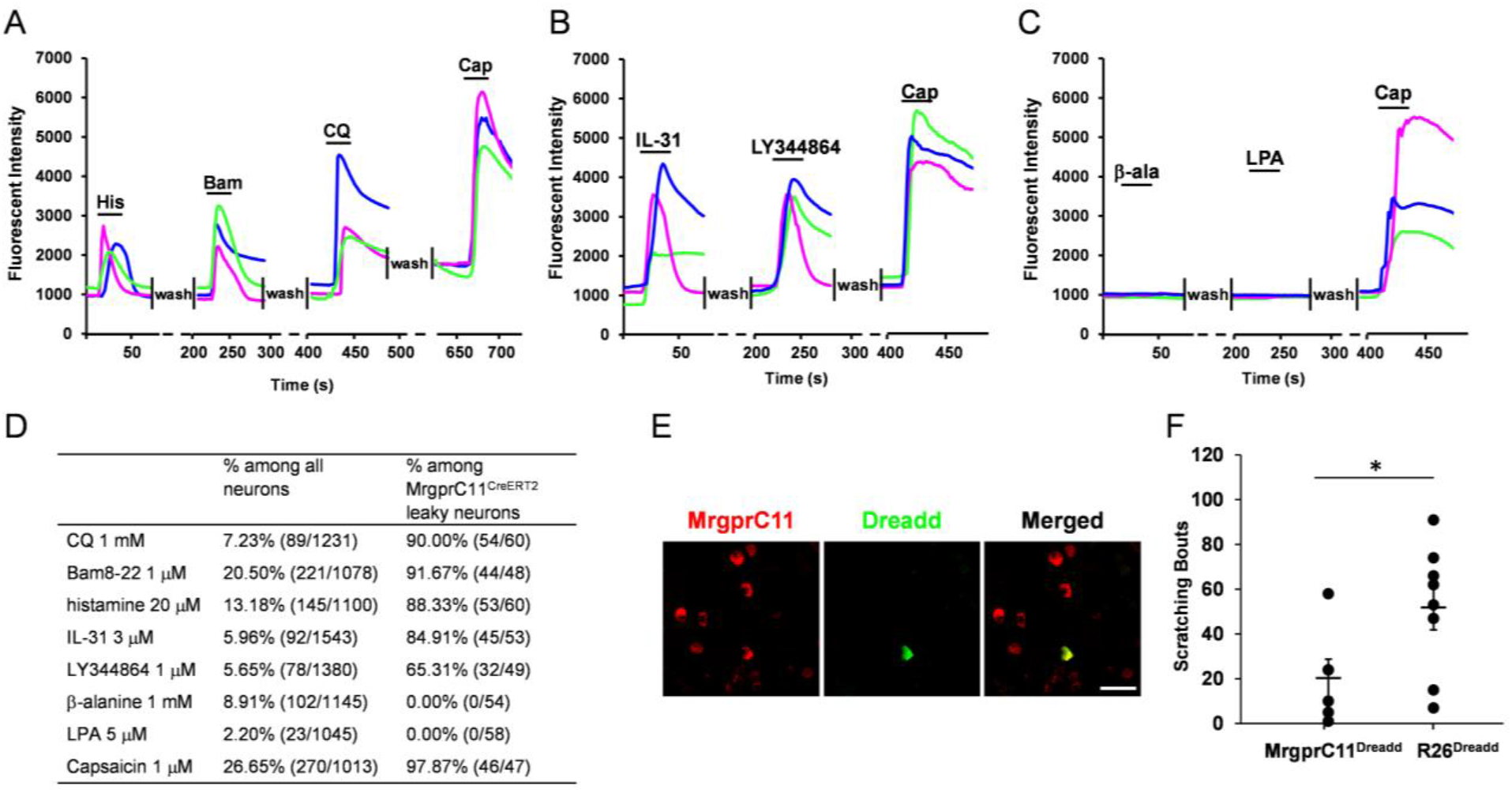
MrgprC11^CreERT2^ leaky neurons represent itch-sensing neurons. (A-C) Representative traces evoked by different pruritogens in cultured DRG sensory neurons isolated from the *MrgprC11^tdtomato^; Pirt^GCaMP3^* mice. (D) Table showing the concentration of pruritogens tested (left column), the percentages of responding neurons among all cells (middle column), and the percentages of responding neurons among MrgprC11^CreERT2^ leaky cells (right column). Total numbers of neurons analyzed are shown in parenthesis. N = 3 mice for each pruritogen. (E) Double immunostaining of lumbar DRG sections collected from *MrgprC11^DREADD^* mice using MrgprC11 and GFP antibodies. The hM3Dq transgene in *ROSA26^DREADD^* mice was fused with mCitrine that can be detected by GFP antibody to examine the expression of DREADD in MrgprC11^+^ neurons. Scale bars = 50 μm. (F) Subcutaneous injection of CNO (10 mM) into the nape of the neck induced significant scratching behavior in *MrgprC11^DREADD^* mice compared to *R26^DREADD^* mice. N = 8 for the *MrgprC11^DREADD^* group and N = 6 for the R26^*DREADD*^ group. *P < 0.05. Two-tailed unpaired Student’s t test. Error bars represent s.e.m.

We then employed a chemogenetic approach to examine if activation of MrgprC11^CreERT2^ leaky cells can induce itch behavior in mice. We generated *MrgprC11^DREADD^* mice by crossing our *MrgprC11^CreERT2^* line with the *Rosa26^DREADD^* line. Immunohistochemistry analysis confirmed the expression of Dreadd in MrgprC11^+^ neurons (Figure 3E). Subcutaneous injection of CNO into the nape of the neck skin induced robust scratching behaviors in *MrgprC11^DREADD^* mice, but not in *Rosa26^DREADD^* control mice (Figure 3F). This data indicates that the activation of MrgprC11^CreERT2^ leaky neurons induces itch sensation. In summary, these data suggest that MrgprC11^CreERT2^ leaky neurons, which constitute approximately 1% of DRG sensory neurons, represent itch-sensing neurons Our newly generated *MrgprC11^CreERT2^* line provides a genetic tool for sparse genetic tracing to investigate the morphological features of itch-sensing neurons.

### Itch-sensing skin arbors are characterized by free endings with extensive axonal branching in the superficial epidermis and large receptive fields

We crossed the *MrgprC11^CreERT2^* mice with Cre-dependent *Rosa26^IAP^* line to express the axonal tracer PLAP (human placental alkaline phosphatase) in MrgprC11^CreERT2^ leaky cells. We observed 41-102 PLAP^+^ neurons/DRG (53.4 ±3.6 neurons/cervical DRG, 64.2 ±3.4 neurons/thoracic DRG and 79 ±3.6 neurons/lumbar DRG) (Figure 4A). These sensory arbors cover ~80% of the skin surface area. The arbor density increased proximally and in the upper part of the body. The majority of arbors are overlapping, suggesting that itch-sensing arbors do not tile the skin (Figure 4B). Arbors were observed in the distal limbs, including the forepaws and hindpaws, but only in the dorsal skin of the paws, not in the glabrous plantar skin (Figure 4C). The morphology of MrgprC11^+^ arbors in the skin is consistent among animals, and are typical arbors with “free endings” featuring extensive axonal branching uniformly covering the whole area (Figure 4D). These features indicate that the axon terminals are free nerve endings in the superficial epidermis. Circumferential-like endings associated with hair follicles were occasionally observed (Figure 4D) in approximately 10% of the arbors. This innervation pattern was confirmed by skin sections from tamoxifen-treated *MrgprC11^tdTomato^* mice. tdTomato^+^ skin nerves penetrate to the most superficial skin layer (Figure 4F), a feature typical of itch-sensing nerves (Han et al., 2013). We occasionally observed circumferential endings with tdTomato^+^ nerves wrapping around hairy follicles (Figure 4F).

**Figure 4.**
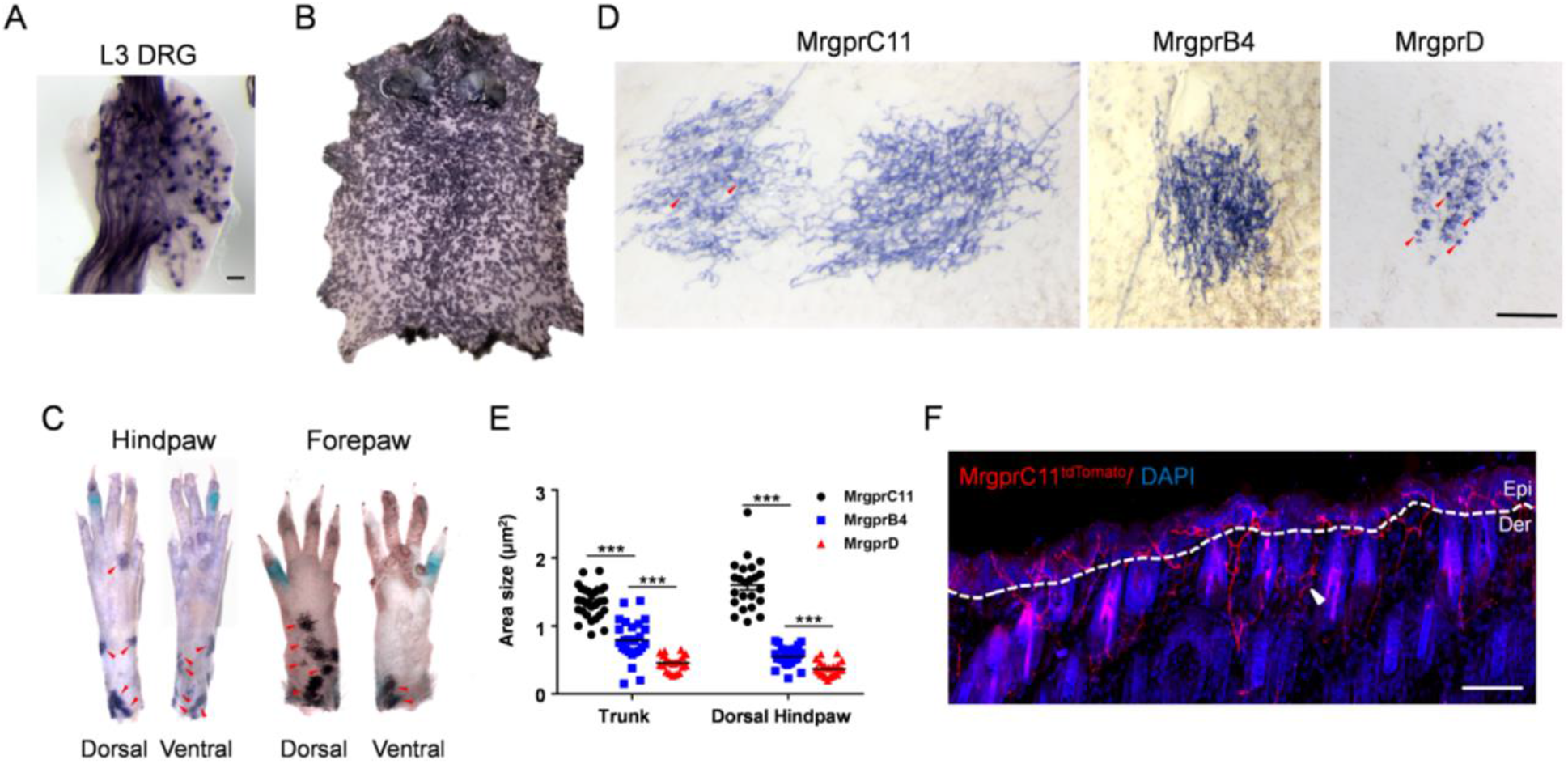
Itch-sensing skin arbors exhibit free nerve endings in the superficial epidermis, dense axon density, and large receptive fields. (A) Whole-mount PLAP histochemistry staining of a lumbar level 3 (L3) DRG from *MrgprC11^IAP^* mouse. Approximately 1% of DRG sensory neurons were labeled by the axonal tracer PLAP. (B) Whole-mount PLAP histochemistry of whole body skin showing the distribution of MrgprC11^+^ skin arbor. (C) Whole-mount PLAP histochemistry showing the distribution of MrgprC11^+^ skin arbor on distal limbs. Arbors, indicated by the red arrows, were observed in the dorsal hairy skin of the paws, not in the ventral plantar glabrous skin. Multiple high magnification images were stitched together to show the whole paws. (D) High magnification images showing single skin arbor morphology using *MrgprC11^IAP^, MrgprB4^PLAP^, and MrgprD^IAP^* mice. Circumferential-like endings associated with hair follicles were observed frequently in almost every MrgprD^+^ arbors and occasionally in MrgprC11^+^ arbors. Some of them are indicated with red arrowheads. Two neighboring MrgprC11^+^ arbors were shown with one of them exhibiting several circumferential-like endings. Scale bars = 500 μm. (E) Areas of the three subtypes of nociceptive skin arbors in the trunk and dorsal hindpaw. Data present as mean ± s.e.m. (F) Skin cross-section from the trunk area of tamoxifen-treated *MrgprC11^tdTomato^* mouse. MrgprC11^+^ nerves are free nerve endings penetrating into the superficial layers of the epidermis. Nerves wrapping around the hair follicles, indicated by the white arrowhead, were occasionally observed. Scale bars = 50 μm.

### Itch-sensing skin arbors exhibit large receptive fields

We compared the morphology of itch-sensing skin arbors with MrgprD^CreERT2^ leaky neurons and MrgprB4^+^ neurons (Figure 4D). MrgprD^CreERT2^ leaky neurons represent non-peptidergic nociceptors whose arbor morphology was revealed by the background recombination of *MrgprD^CreERT2^* line (Olson et al., 2017; Olson and Luo, 2018). Consistent with the previous report, we found that these arbors have “bushy ending” morphology, featuring dense and finely branched processes and clustered terminal neurites. Their axons show both free nerve endings in the epidermis and frequent circumferential endings (Figure 4D). MrgprB4^+^ neurons, which constitute about 2% of DRG sensory neurons, are mechanosensory C-fibre mediating pleasant stroking of hairy skin. Their skin arbors also exhibit free ending morphology visualized using *MrgprB4^PLAP^* knockin mouse line (Figure 4D) (Liu et al., 2007). Arbor areas were significantly different, with MrgprC11^CreERT2^ leaky neurons showing the largest arbor area and MrgprD^CreERT2^ leaky neurons showing the smallest. Only non-overlapping MrgprC11^+^ arbors in the trunk skin and dorsal hindpaw skin were selected for quantification. The size difference is more prominent in the dorsal hindpaw skin with the size of MrgprC11^+^ arbors triple the size of the other two subtypes (1.60 ± 0.08 mm^2^ for MrgprC11^CreERT2^ leaky neurons, 0.55 ± 0.03 mm^2^ for MrgprB4^+^ neurons, 0.37 ± 0.02 mm^2^ for MrgprD^CreERT2^ leaky neurons) (Figure 4E). In summary, these results demonstrate the distinct terminal arborizations of three C-type cutaneous afferents. Itch-sensing skin arbors, revealed for the first time using MrgprC11^CreERT2^ leaky neurons, are characterized by free endings with extensive axonal branching in the superficial epidermis and large receptive fields.

### MrgprD^CreERT2^ leaky neurons exhibit limited expression of itch receptors

Since MrgprD mediates β-alanine-induced itch sensation, we asked if MrgprD^CreERT2^ leaky neurons also represent a subset of itch-sensing neurons. We examined the molecular feature of MrgprD^CreERT2^ leaky neurons from *MrgprE^tdtomato^* mice (Figure 5A-H). All labeled neurons expressed MrgprD and most also expressed Lpar3 (83.9%), which is highly enriched in MrgprD^+^ neurons (Figure 5A-B). However, none of them co-expressed with the other examined itch receptors including MrgprA3, MrgprC11, Il31ra, and Htrf1. They do not express Nppb and very few co-expressed the histamine receptor Hrh1 (5.33%) (Figure 4C-H). Calcium imaging with sensory neurons isolated from *MrgprD^tdTomato;^ Pirt^GCaMP3^* mice showed that a small percentage of MrgprD^CreERT2^ leaky neurons responded to β-alanine (8.6%). This is not a surprise since only half of MrgprD^+^ neurons were reported to respond to β-alanine (Liu et al., 2012). 14.6% of the labeled neurons responded to LPA (Figure 5I-J). Only a few (0-2, 0% - 4.6%) tdTomato^+^ cells responded to other chemicals including chloroquine, Bam8-22, histamine, Il-31, LY344864, and capsaicin (Figure 5J). Since MrgprD^CreERT2^ leaky neurons exhibit limited expression of itch receptors and weak response to pruritogens, we do not have strong evidence to suggest that MrgprD^CreERT2^ leaky neurons are itch-sensing neurons.

**Figure 5.**
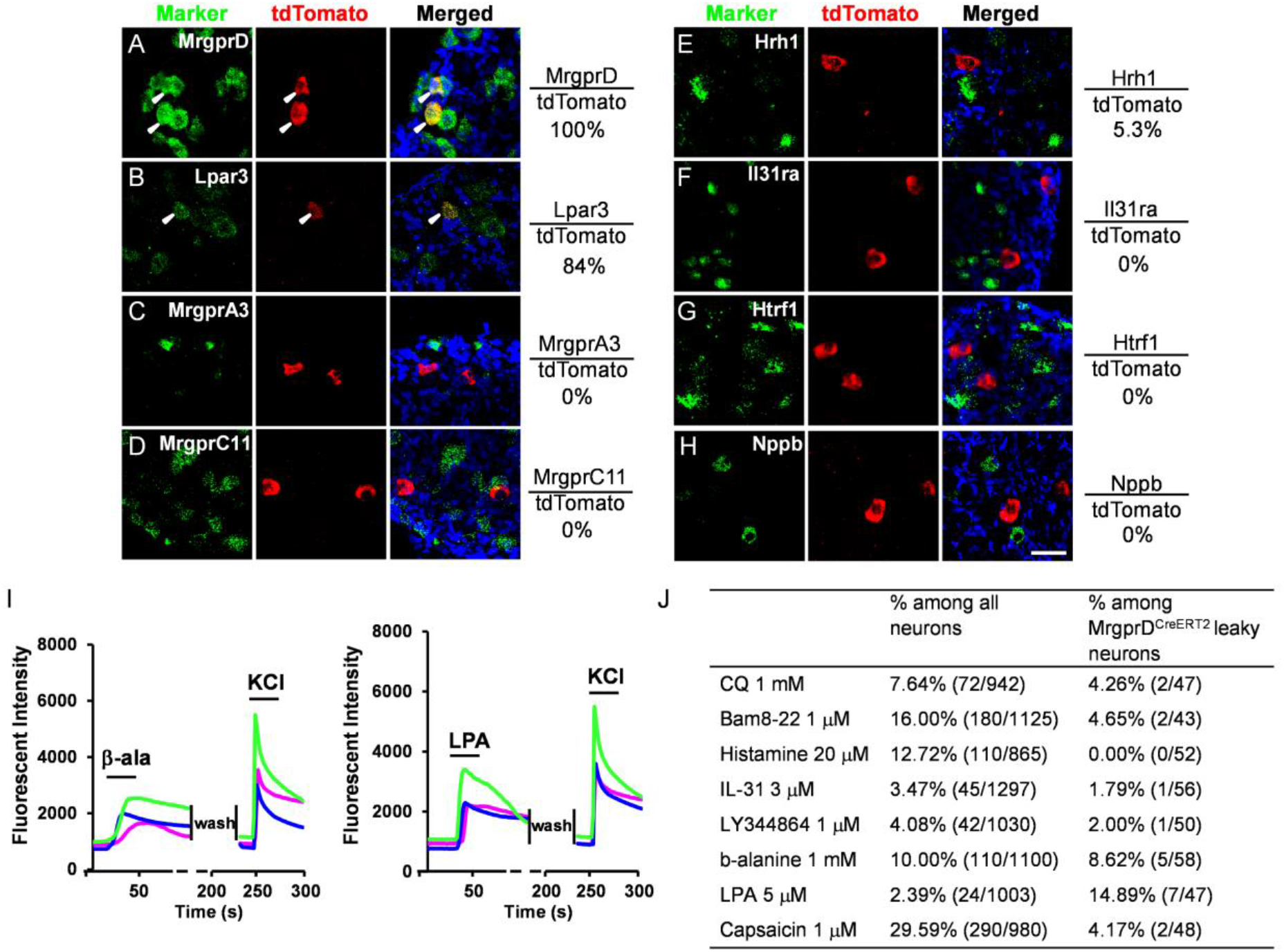
MrgprD^CreERT2^ leaky neurons express limited itch receptors. (A-H) RNAscope fluorescent *in situ* hybridization detecting itch-related molecules using lumbar DRG sections from *MrgprD^tdTomato^* mice. Overlap quantifications were shown on the right of each panel (% of tdTomato^+^ cells that express markers). Arrowheads point to overlapping cells. Scale bars = 50 μm. N = 3 mice for each marker. (I) Representative traces evoked by different pruritogens in cultured DRG sensory neurons isolated from the *MrgprD^tdtomato^; Pirt^GCaMP3^* mice. A small percentage of MrgprD^CreERT2^ leaky neurons responded to β-alanine and LPA. All cells were subjected to KCl (50 mM) treatment to verify their neuronal identity. (J) Table showing the concentration of pruritogens tested (left column), the percentages of responding neurons among all cells (middle column), and the percentages of responding neurons among MrgprD^CreER^ leaky cells (right column). Total numbers of neurons analyzed were shown in parenthesis. N = 2 mice for each pruritogen.

## DISCUSSION

We generated a novel mouse line, *MrgprC11^CreERT2^*, which labels a subset of peptidergic itch-sensing neurons responding to multiple pruritogens. Using sparse genetic tracing, we revealed the morphological features of the itch-sensing arbors in the skin.

MrgprA3^+^ neurons and Nppb^+^ neurons are two distinct itch-sensing neuronal populations with minimal overlap (Solinski et al., 2019). A common feature of itch-sensing neurons is that they are enriched with multiple itch receptors (Sharma et al., 2020; Solinski et al., 2019; Usoskin et al., 2015; Xing et al., 2020). MrgprA3^+^ neurons express histamine receptors and two other identified itch receptors in Mrgpr family: MrgprA1 and MrgprC11. Nppb^+^ neurons are highly enriched with IL-31 receptors, Cysltr2, Htr1f, and S1pr1. The majority of MrgprC11^CreERT2^ leaky neurons we identified using sparse genetic labeling express markers for both itch-sensing populations including MrgprA3, MrgprC11, Hrh1, Nppb, IL31ra, Cysltr2, and Htr1f. Consistently, calcium imaging analysis suggests that most MrgprC11^CreERT2^ leaky neurons respond to the corresponding pruritogens. This molecular feature suggests that these neurons likely represent the overlapping piece of the two itch-sensing neuronal populations. Indeed, activation of MrgprC11^CreERT2^ leaky neurons using a chemogenetic approach induced robust scratching behaviors, demonstrating that they are itch-sensing neurons. Therefore, the *MrgprC11^CreERT2^* transgenic line we generated serves as a great genetic tool for the morphological characterization of itch-sensing neurons.

Sensory arbor morphology evolved to properly serve the physiological function of neurons. Our comparison of the peripheral arbors of MrgprC11^CreERT2^ leaky neurons, MrgprB4^+^ neurons, and MrgprD^CreERT2^ leaky neurons highlights the diversity of the cutaneous C-type afferents. MrgprC11^+^ itch-sensing arbors exhibit extensive axonal branching and free nerve endings in the superficial epidermis, suggesting that itch-sensing neurons are highly sensitive to pruritic stimuli on the surface of the skin. Circumferential-like endings associated with hair follicles were only occasionally observed in a small percentage of arbors (~10%), therefore they do not seem to be good candidates for mediating itch induced by hair vibration, a form of mechanically evoked itch (Fukuoka et al., 2013).

Sensory arbor area represents the receptive field size of sensory neurons. More than 10 morphologically distinct cutaneous arbor types have been characterized using sparse genetic labeling (Abraira and Ginty, 2013; Badea et al., 2012; Bai et al., 2015; Kuehn et al., 2019; Li and Ginty, 2014; Li et al., 2011; Rutlin et al., 2014; Wu et al., 2012) and the majority of them are large-diameter Aβ or Aδ fibers that are associated with specialized skin structure such as hairy follicles and merkel cells. Among them, the Aβ field LTMR innervating the hairy follicles exhibit a large receptive field (~ 3 mm^2^) to detect gentle stroking across large fields of hairy skin (Bai et al., 2015). To our knowledge, four subtypes of free nerve endings have been identified: three subtypes reported here and by previous studies (Liu et al., 2007; Olson et al., 2017) and large area free ending (LA-FE) identified by random sparse labeling of *NFL^CreERT2^* line (Wu et al., 2012). LA-FE arbors were named based on the morphological features and their function is unknown. They have the largest receptive fields among all reported (10-30 mm^2^) with dense axonal branches uniformly distributed within the arbor (Wu et al., 2012). The area of itch-sensing neurons (1.60 ± 0.08 mm^2^ in the dorsal hindpaw) is larger than the other two free ending arbors (MrgprB4^+^ and MrgprD^+^) and most of the non-free ending arbors. IT is worth noting that the area of itch-sensing arbors may be slightly underestimated as only isolated arbors in the thoracic and lumbar regions were selected for analysis, whose sizes tend to be smaller than the overlapping ones. Interestingly, a study from human subjects suggests that itch-sensing neurons have large receptive fields. Schmelz et al. showed that a group of mechanically insensitive and histamine sensitive C-fiber, presumably itch-sensing, have much larger innervation territories (up to 45-88 mm diameter) than the histamine insensitive C fibers examined in human skin (Schmelz et al., 1997). This might explain why itch sensation is normally perceived as rising from a diffused large area instead of a focused spot.

## MATERIAL AND METHODS

### Animals

All experiments were performed with approval from the Georgia Institute of Technology Animal Use and Care Committee. *MrgprC11^CreERT2^* BAC transgenic line, *MrgprB4^PLAP^* knockin line (Liu et al., 2007), and *Pirt^GCamp3^* knockin line (Kim et al., 2014) were generously provided by Dr. Xinzhong Dong at the Johns Hopkins University. *MrgprC11^CreERT2^* BAC transgenic mouse line was generated by the Gene Targeting & Transgenic Facility at Janelia Farm. *MrgprD^CreERT2^* (Stock No: 031286), *ROSA26^tdTomato^* (Stock No: 007914), *ROSA26^IAP^* (Stock No: 009253), and *ROSA26^DREADD^* (Stock No: 026220) mouse lines were purchased from Jackson Laboratory. All the mice used had been backcrossed to C57BL/6 mice for at least ten generations. All the animals were housed in the vivarium with a 12-h light/dark cycle, and all the behavioral tests were performed from 9 a.m. to 1 p.m. in the light cycle. Mice were housed in groups, with a maximum of five per cage with food and water *ad libitum*.

### PLAP whole-mount histochemistry staining

The procedure was performed as previously described (Liu et al., 2007). The tissues were cleared in BABB (Benzyl Alcohol and Benzyl Benzoate, 1:2 mixed together) before imaging on a ZEISS SteREO Discovery V12 stereomicroscope with a color camera. Two to five DRGs from one animal were used to quantify the number of PLAP^+^ cells in DRGs collected from different body areas. For the quantification of the size of both skin arbors and central arbors, mice with similar age (P28 to P35) and body weight were used to avoid variation caused by animal size.

### Immunohistochemistry staining and RNAscope *In situ* Hybridization

Immunohistochemistry staining was performed as previously described (Han et al., 2013). The following antibodies were used: rabbit anti-MrgprC11 (custom made by Proteintech groups, validated by previous study, 1:500) (Han et al., 2018), rabbit anti-TRPV1 (Novus, NBP120-3487, 1:500), rat anti-Substance P (Millipore, MAB356, 1:500), goat anti-CGRP (BIO-RAD, 1720-9007, 1:500), conjugated IB4-Alexa 488 (Thermo Fisher, I-21411, 1:200), rabbit anti-P2X3 (Millipore, AB5898, 1:500), chicken anti-NF200 (Aves Labs, NFH877982, 1:500), rabbit anti-HA to detect hM3Dq expression (Cell Signaling Technologies, 3724S, 1:500), donkey anti-rabbit Alexa Fluor 488 (Thermo Fisher, A11056), goat anti-chicken Alexa Fluor 488 (Thermo Fisher, A21127), donkey anti-goat Alexa Fluor 488 (Thermo Fisher, A21206), donkey anti-goat Alexa Fluor 546 (Thermo Fisher, A31572), and donkey anti-rat Alexa Fluor 488 (Thermo Fisher, A11039). All secondary antibodies were used at 1:500 dilution. To visualize tdTomato^+^ sensory nerves in the skin, *MrgprC11^tdTomato^* mice were treated daily via oral gavage (22g x 25mm, FTP-22-25, Instech Laboratories) with 100 mg/kg of tamoxifen (T5648, Sigma) dissolved in sunflower seed oil (S5007, Sigma) at P14-P18. Cross skin sections were collected and tdTomato^+^ nerves were directly imaged with Confocal Microscopy.

RNAscope *in situ* hybridization was performed using the RNAscope fluorescent multiplex kit (ACD Cat#320850) according to the manufacturer’s instructions. The following probes were used: tdTomato (317041), MrgprA3 (548161), MrgprC11 (488771), MrgprD (417921), Hrh1 (491141), Il31ra (418411), Htrf1 (315881), Nppb (425021), Nmb (459931), Lpar3 (432591).

Quantification results for every marker were collected from 4-12 lumbar DRGs dissected from 2-3 adult mice. The number of neurons quantified for all analyses are indicated in the figures.

### Behavioral tests

The experiments were performed as previously described (Xing et al., 2020). Briefly, Clozapine N-oxide (CNO 16882, Cayman, 10 mg/kg, 100 μl) were injected subcutaneously into the nape of the neck. Behavioral responses were video recorded and analyzed by experimenters blind to genotype.

### Calcium Imaging

Calcium imaging was performed with sensory neurons collected from *MrgprC11^tdTomato^; Pirt^GCaMP3^* mice or *MrgprD^tdTomato^; Pirt^GCaMP3^* mice. Three *MrgprC11^tdTomato^; Pirt^GCaMP3^* mice and two *MrgprD^tdTomato^; Pirt^GCaMP3^* mice were used for calcium imaging and quantification. A 20% increase in GCaMP3 fluorescence intensity was set as the threshold to identify responding cells. Chemicals used are listed as follows: chloroquine (Sigma PHR1258), Bam8-22 (custom synthesized by Genscript), histamine (Sigma H7250), IL-31 (R&D 3028-ML-010), LY344864 (Sigma SML0556), β-alanine (Sigma 146064), Capsaicin (Sigma M2028), LPA 18:1 (Sigma 857130).

## AUTHOR CONTRIBUTION

Conceptualization: LH; Formal Analysis: YX, HS, HH, TN, KL, LH; Investigation: YX, HS, HH, TN, KL, LH; Resources: KL; Writing - Original Draft Preparation: YX, LH; Writing - Review and Editing: YX, HS, LH; Supervision: LH; Funding Acquisition: LH.

## CONFLICT OF INTEREST

The authors claim no conflict of interest.

## ACKNOWLEDGMENTS

We thank the Physiological Research laboratory (PRL) at Georgia Institute of Technology for the animal care and services. The work was supported by grants from the US National Institutes of Health (NS087088 and HL141269) and Pfizer Aspire Dermatology Award to L.H.

## Notes

### Competing Interest Statement

The authors have declared no competing interest.

## Reference

Abraira, V.E., and Ginty, D.D. (2013). The sensory neurons of touch. Neuron 79, 618–639.

Badea, T.C., Williams, J., Smallwood, P., Shi, M., Motajo, O., and Nathans, J. (2012). Combinatorial expression of Brn3 transcription factors in somatosensory neurons: genetic and morphologic analysis. J Neurosci 32, 995–1007.

Bai, L., Lehnert, B.P., Liu, J., Neubarth, N.L., Dickendesher, T.L., Nwe, P.H., Cassidy, C., Woodbury, C.J., and Ginty, D.D. (2015). Genetic Identification of an Expansive Mechanoreceptor Sensitive to Skin Stroking. Cell 163, 1783–1795.

Basbaum, A.I., Bautista, D.M., Scherrer, G., and Julius, D. (2009). Cellular and molecular mechanisms of pain. Cell 139, 267–284.

Crawford, L.K., and Caterina, M.J. (2020). Functional Anatomy of the Sensory Nervous System: Updates From the Neuroscience Bench. Toxicologic pathology 48, 174–189.

Fukuoka, M., Miyachi, Y., and Ikoma, A. (2013). Mechanically evoked itch in humans. Pain 154, 897–904.

Ghitani, N., Barik, A., Szczot, M., Thompson, J.H., Li, C., Le Pichon, C.E., Krashes, M.J., and Chesler, A.T. (2017). Specialized Mechanosensory Nociceptors Mediating Rapid Responses to Hair Pull. Neuron 95, 944–954 e944.

Han, L., Limjunyawong, N., Ru, F., Li, Z., Hall, O.J., Steele, H., Zhu, Y., Wilson, J., Mitzner, W., Kollarik, M., et al. (2018). Mrgprs on vagal sensory neurons contribute to bronchoconstriction and airway hyper-responsiveness. Nat Neurosci 21, 324–328.

Han, L., Ma, C., Liu, Q., Weng, H.J., Cui, Y., Tang, Z., Kim, Y., Nie, H., Qu, L., Patel, K.N., et al. (2013). A subpopulation of nociceptors specifically linked to itch. Nat Neurosci 16, 174–182.

Ikoma, A., Steinhoff, M., Stander, S., Yosipovitch, G., and Schmelz, M. (2006). The neurobiology of itch. Nat Rev Neurosci 7, 535–547.

Kim, Y.S., Chu, Y., Han, L., Li, M., Li, Z., LaVinka, P.C., Sun, S., Tang, Z., Park, K., Caterina, M.J., et al. (2014). Central terminal sensitization of TRPV1 by descending serotonergic facilitation modulates chronic pain. Neuron 81, 873–887.

Kuehn, E.D., Meltzer, S., Abraira, V.E., Ho, C.Y., and Ginty, D.D. (2019). Tiling and somatotopic alignment of mammalian low-threshold mechanoreceptors. Proc Natl Acad Sci U S A 116, 9168–9177.

Li, L., and Ginty, D.D. (2014). The structure and organization of lanceolate mechanosensory complexes at mouse hair follicles. Elife 3, e01901.

Li, L., Rutlin, M., Abraira, V.E., Cassidy, C., Kus, L., Gong, S., Jankowski, M.P., Luo, W., Heintz, N., Koerber, H.R., et al. (2011). The functional organization of cutaneous low-threshold mechanosensory neurons. Cell 147, 1615–1627.

Liu, Q., Sikand, P., Ma, C., Tang, Z., Han, L., Li, Z., Sun, S., LaMotte, R.H., and Dong, X. (2012). Mechanisms of itch evoked by beta-alanine. J Neurosci 32, 14532–14537.

Liu, Q., Tang, Z., Surdenikova, L., Kim, S., Patel, K.N., Kim, A., Ru, F., Guan, Y., Weng, H.J., Geng, Y., et al. (2009). Sensory neuron-specific GPCR Mrgprs are itch receptors mediating chloroquine-induced pruritus. Cell 139, 1353–1365.

Liu, Q., Vrontou, S., Rice, F.L., Zylka, M.J., Dong, X., and Anderson, D.J. (2007). Molecular genetic visualization of a rare subset of unmyelinated sensory neurons that may detect gentle touch. Nat Neurosci 10, 946–948.

Liu, Q., Weng, H.J., Patel, K.N., Tang, Z., Bai, H., Steinhoff, M., and Dong, X. (2011). The distinct roles of two GPCRs, MrgprC11 and PAR2, in itch and hyperalgesia. Sci Signal 4, ra45.

Olson, W., Abdus-Saboor, I., Cui, L., Burdge, J., Raabe, T., Ma, M., and Luo, W. (2017). Sparse genetic tracing reveals regionally specific functional organization of mammalian nociceptors. Elife 6.

Olson, W., and Luo, W. (2018). Somatotopic organization of central arbors from nociceptive afferents develops independently of their intact peripheral target innervation. The Journal of comparative neurology 526, 3058–3065.

Reddy, V.B., Sun, S., Azimi, E., Elmariah, S.B., Dong, X., and Lerner, E.A. (2015). Redefining the concept of protease-activated receptors: cathepsin S evokes itch via activation of Mrgprs. Nat Commun 6, 7864.

Rutlin, M., Ho, C.Y., Abraira, V.E., Cassidy, C., Bai, L., Woodbury, C.J., and Ginty, D.D. (2014). The cellular and molecular basis of direction selectivity of Adelta-LTMRs. Cell 159, 1640–1651.

Schmelz, M., Schmidt, R., Bickel, A., Handwerker, H.O., and Torebjork, H.E. (1997). Specific C-receptors for itch in human skin. J Neurosci 17, 8003–8008.

Sharma, N., Flaherty, K., Lezgiyeva, K., Wagner, D.E., Klein, A.M., and Ginty, D.D. (2020). The emergence of transcriptional identity in somatosensory neurons. Nature 577, 392–398.

Solinski, H.J., Kriegbaum, M.C., Tseng, P.Y., Earnest, T.W., Gu, X., Barik, A., Chesler, A.T., and Hoon, M.A. (2019). Nppb Neurons Are Sensors of Mast Cell-Induced Itch. Cell Rep 26, 3561-3573 e3564.

Usoskin, D., Furlan, A., Islam, S., Abdo, H., Lonnerberg, P., Lou, D., Hjerling-Leffler, J., Haeggstrom, J., Kharchenko, O., Kharchenko, P.V., et al. (2015). Unbiased classification of sensory neuron types by large-scale single-cell RNA sequencing. Nat Neurosci 18, 145–153.

Wan, L., Jin, H., Liu, X.Y., Jeffry, J., Barry, D.M., Shen, K.F., Peng, J.H., Liu, X.T., Jin, J.H., Sun, Y., et al. (2017). Distinct roles of NMB and GRP in itch transmission. Sci Rep 7, 15466.

Wu, H., Williams, J., and Nathans, J. (2012). Morphologic diversity of cutaneous sensory afferents revealed by genetically directed sparse labeling. Elife 1, e00181.

Xing, Y., Chen, J., Hilley, H., Steele, H., Yang, J., and Han, L. (2020). Molecular signature of pruriceptive MrgprA3(+) neurons. J Invest Dermatol.

Zylka, M.J., Rice, F.L., and Anderson, D.J. (2005). Topographically distinct epidermal nociceptive circuits revealed by axonal tracers targeted to Mrgprd. Neuron 45, 17–25.

